# Progressive circuit hyperexcitability in mouse neocortical slice cultures with increasing duration of activity silencing

**DOI:** 10.1101/2023.07.07.548151

**Authors:** Derek L. Wise, Samuel B. Greene, Yasmin Escobedo-Lozoya, Stephen D. Van Hooser, Sacha B. Nelson

## Abstract

Forebrain neurons deprived of activity become hyperactive when activity is restored. Rebound activity has been linked to spontaneous seizures *in vivo* following prolonged activity blockade. Here we measured the time course of rebound activity and the contributing circuit mechanisms using calcium imaging, synaptic staining and whole cell patch clamp in organotypic slice cultures of mouse neocortex. Calcium imaging revealed hypersynchronous activity increasing in intensity with longer periods of deprivation. While activity partially recovered three days after slices were released from five days of deprivation, they were less able to recover after ten days of deprivation. However, even after the longer period of deprivation, activity patterns eventually returned to baseline levels. The degree of deprivation-induced rebound was age-dependent, with the greatest effects occurring when silencing began in the second week. Pharmacological blockade of NMDA receptors indicated that hypersynchronous rebound activity did not require Hebbian plasticity evoked. In single neuron recordings, input resistance roughly doubled with a concomitant increase in intrinsic excitability. Synaptic imaging of pre- and postsynaptic proteins revealed dramatic reductions in the number of presumptive synapses with a larger effect on inhibitory than seen in excitatory synapses. Putative excitatory synapses colocalizing PSD-95 and Bassoon declined by 39% and 56% following five and ten days of deprivation, but presumptive inhibitory synapses colocalizing gephyrin and VGAT declined by 55% and 73% respectively. The results suggest that with prolonged deprivation, a progressive reduction in synapse number is accompanied by a shift in the balance between excitation and inhibition and increased cellular excitability.

**SIGNIFICANCE STATEMENT:** When cortical activity is silenced during development, lifelong seizures often result. Here we explored whether these seizures result from overcompensation of homeostatic recovery mechanisms. Prior work showed that neurons briefly deprived of the ability to fire action potentials compensate by becoming more excitable, increasing synaptic drive and intrinsic excitability. We found that prolonged silencing of cortex produced a profound loss of synapse density, especially for inhibitory synapses, pointing to a circuit unable to maintain excitatory/inhibitory balance. These results show that homeostatic responses, which are normally restorative, can result in maladaptive circuit configurations when brought to extremes.

## INTRODUCTION

During development, forebrain circuits respond homeostatically to changes in activity, decreasing circuit excitability when activity increases, (Kilman et al., 2002; Swanwick et al., 2006) and increasing circuit excitability to counteract decreased activity (Desai et al., 1999; Turrigiano et al., 1998). However, this normally beneficial homeostatic plasticity can be maladaptive if compensatory mechanisms overshoot normal activity levels (Nelson and Valakh, 2015). Silencing of neuronal activity during development is sufficient to produce life-long epilepsy (Galvan et al., 2000; Lee et al., 2008): a protracted activity blockade (10-14 days) that was delivered during the second postnatal week induced a lasting susceptibility to seizures.

Blocking activity later in development did not cause seizures, suggesting a particular developmental vulnerability. This developmental stage is also a time when homeostatic regulation of circuit excitability is known to be stronger than it is later in development (Desai et al., 2002).

Previous work has examined homeostatic responses to short-term (4-48 hour) reductions of neuronal activity. These include changes in postsynaptic glutamate receptor trafficking (Ehlers, 2000; Gainey et al., 2009; O’Brien et al., 1998), changes in presynaptic glutamate release (Echegoyen et al., 2007; Frank et al., 2006; Murthy et al., 2001), changes in the pre and postsynaptic strength of inhibitory synapses (Bartley et al., 2008; Chattopadhyaya et al., 2004; Kilman et al., 2002; Marty et al., 2000), and changes in cellular excitability (Desai et al., 1999; LeMasson et al., 1993).

By contrast, far less is known about the consequences of longer periods of activity blockade, or about how activity recovers once it is restored. Prolonged activity deprivation increases susceptibility to seizures as described above (Galvan et al., 2000; Lee et al., 2008) but the underlying mechanisms are not known. Persistent hyperactivity following removal of activity blockade could reflect altered cellular excitability and/or changes in the strength and number of excitatory and inhibitory synapses brought about as part of the initial homeostatic response. In addition, other forms of plasticity, such as Hebbian Long-term Potentiation (LTP) or Depression (LTD) could further alter synaptic strength once network activity resumes.

In order to learn more about the kinetics of this hypersynchronous activity overshoot, we blocked neuronal firing with the sodium channel blocker tetrodotoxin (TTX) in organotypic cortical slice cultures and then monitored network activity with calcium imaging at various times after removal of TTX. To assess the impact of Hebbian plasticity, we compared recovery in the presence and absence of the NMDA receptor blocker APV. To begin to tease apart some of the cellular and synaptic factors that contribute to abnormal network activity following protracted activity blockade, we then measured cellular excitability in whole cell recordings and imaged presumptive excitatory and inhibitory synapses with super resolution confocal microscopy. The results reveal complex progressive and multifactorial changes in these cortical networks that likely contribute to their abnormal hypersynchronous activity and that may contribute to seizures following activity blockade *in vivo*.

## METHODS

### Organotypic cortical slice culture

7-day old wild-type C57-B6J mice were anesthetized with 40 μL of a standard ketamine (20 mg/mL), xylazine (2.5 mg/mL), and acepromazine (0.5 mg/mL) mixture administered intraperitoneally, then decapitated and dissected for resectioning (Stoppini et al., 1991). After being embedded in agarose within a cylindrical chamber that slightly narrows at its opening, the brain was cut using a compresstome (Precisionary Instruments, Greenville, NC) in filtered ice-chilled artificial cerebrospinal fluid (ACSF; 126 mM NaCl, 25 mM NaHCO3, 3 mM KCl, 1 mM NaH2PO4 H20, 25 mM dextrose, 2 mM CaCl and 2 mM MgCl2, 315-319 mOsm) to 300 μm thickness and grown on Millipore Millicell inserts (Millipore Sigma, Burlington, Massachusetts) in 6-well dishes over SCM media (1x MEM (Millipore-Sigma), 1x GLUTAMAX (Gibco Thermo-Fisher Scientific), 1 mM CaCl2, 2 mM MgSO4, 12.9 mM dextrose, 0.08% ascorbic acid, 18 mM NaHC03, 35 mM HEPES (pH 7.5), 1 μg/mL insulin and 20% Horse Serum (heat inactivated, Millipore-Sigma, Burlington, Massachusetts), pH 7.45 and 305 mOsm). A 1,000 units/mL PenStrep (Gibco Thermo-Fisher Scientific, Waltham MA) & 50 μg/mL gentamicin (Millipore-Sigma, Burlington MA) antibiotic mixture was applied for the first 24 hours in a 35°C incubator with 5% CO2 in the atmosphere. After changing into non-antibiotic media, media changes were performed every 2 days for the course of experiments. One day after dissection, a 1 μL dot of AAV9/hSyn/GCaMP6f virus (Addgene, Watertown, MA) diluted 1-5x from commercial titer (2.8x10^13^ to ∼1x10^12^ units/mL) in ACSF, was applied to the surface of the slice just inferior to the cortex.

Cultures were kept active for 15 days in total for most of our experiments (starting at equivalent postnatal day (EP) 7 and stretching to EP25. For one experiment, experiments continued until EP32. During the first 5 days in vitro (DIV), all cultures recovered from dissection. At 5 DIV, tetrodotoxin (TTX) was dissolved in warmed media at 500 nM concentration and applied to some cultures like any other media change. Others continued with no drug treatment, and others had TTX added to the media beginning at 10 DIV. Over the course of these experiments, several informal checks confirmed that full silencing of these cultures occurred within 5-10 minutes after TTX application, and that the drug was still effective at suppressing calcium activity after 48 hours. Recovery experiments took place following a triplicate media change into fresh non-drug media and three days of time to allow for cultures to equilibrate.

### Calcium imaging

Layer 5 of somatosensory cortex was imaged at 33 frames per second for 10 minutes on a Leica DMI 6000B microscope (Leica Microsystems, Inc., Buffalo Grove, IL) with an Andor CSU W1 confocal spinning disc unit, using an Andor Neo sCMOS camera operated by Andor IQ3 (Andor Technology PLC, Belfast, N. Ireland) under perfusion with 35-37°C oxygenated ACSF (the same solution as for dissection). Around half a mm square was imaged with a 10x objective (Zeiss, Oberkochen, Germany), visualizing many hundreds of cells. Recordings of slices that had been treated with TTX were conducted in ACSF that was free of drug, allowing for cellular activity.

### Calcium analysis

Videos were processed in a custom MATLAB suite (Li et al., 2008; RRID:SCR_023369). Ellipsoidal cellular ROIs of neuron somata were selected by hand using a standard deviation projection of the videos, then their average intensity was calculated at each frame and recorded as a raw time trace. These traces were normalized by a standard ΔF/F formula, (value-baseline)/baseline, where the baseline was defined as the average of timepoints where a cell had a low coefficient of variation for the following 50 frames. Noisy or especially dim data was discarded, both for individual cells and also for slices that had less than 15 cells that fit our criteria.

Periods of activity, termed ‘up states’ (Cossart et al., 2003; Johnson and Buonomano, 2007), were detected with knowledge of the slope of the mean trace of all cells’ activity. When there was a significant positive deflection in slope (> 0.15 ΔF/F) in the trace, this was taken to indicate the presence of firing activity in our population. The threshold was experimentally determined to well separate noise from events under our common noise conditions. Up states persisted until no new influx of calcium was observed beyond a user-defined refractory period.

Up states were counted and then characterized in several ways. The correlation coefficient between each cell pair was determined during each up state and averaged across all cell pairs and all up states (synchrony). We also measured the correlation coefficient between the activity of a cell in one up state to its activity in each other up state (stereotypy), using the first three seconds of each firing period. Duration was defined as the time between the first and last frame when a significant positive slope was observed, meaning that the period at the end of each up state where calcium signal exponentially degrades was not considered to be a part of the up state.

We did not attempt to quantify up state amplitudes using calcium imaging because this slow modality provides poor information about the number of action potentials occurring within a short synchronous burst since many factors, such as GCaMP6f expression levels, contribute to overall amplitude differences across experiments and between cells within an experiment.

### Electrophysiology

Intrinsic excitability was measured from current clamp recordings of visually identified layer 5 pyramidal cells in primary somatosensory cortex in response to a series of 1 s long current pulses from a resting membrane potential of -70mV. Slice cultures were taken from the incubator directly to the electrophysiology rig on a cut-out section of insert and allowed to rest for 20 minutes at 35-37°C under oxygenated ACSF (the same as from dissection) before recording began. Glass recording pipettes (3-5 MΩ resistance) were filled with internal solution (in mM: 100 K-gluconate, 20 KCl, 10 HEPES, 4 Mg-ATP, 0.3 Na-GTP, 10 Na-phosphocreatine, and 0.05% biocytin). Recordings were made using an AxoPatch 200B amplifier (Axon Instruments, Foster City CA), filtered at 10 kHz without correction for liquid junction potentials and digitized at 20 kHz using a National Instruments data acquisition board in a Dell Computer using custom Igor Pro software (WaveMetrics, Lake Oswego, OR). Further analysis was performed in Igor, MATLAB (The Mathworks, Natick, MA) or R.

### Synaptic imaging

Slice cultures were fixed with 3% glyoxal for 30 minutes at 4°C and then 30 minutes at room temperature (Richter et al., 2018). Fixed slices were quenched in ammonium chloride/glycine solution (10 mM each) for 5 minutes, then washed 3 times into PBS and stored at 4°C until staining in PBS containing 1 μM sodium azide.

For staining, single-hemisphere slices were removed from culturing membranes and first exposed to CUBIC 1 solution overnight at 37°C on a shaker (Susaki et al., 2015). These slices were then treated with the DeepLabel Staining system (Logos Bio, Gyeonggi-do South Korea) suitable for thick samples, using a version of the DeepLabel protocol featuring overnight washes in each proprietary solution. Primary antibodies, for our excitatory synaptic stain, were anti-PSD-95 (mouse, Synaptic Systems 124 011, Goettingen Germany), and anti-Bassoon (rabbit, Synaptic Systems 141 003). Secondary antibodies were goat anti-mouse Alexa Fluor Plus 555 (Invitrogen A32727, Carlsbad CA), and goat anti-rabbit Alexa Fluor Plus 488 (Invitrogen A32732). All antibodies here were diluted 1:2000 for use. In the final wash, DAPI (Thermo Fisher Scientific 62248, Waltham MA) was added at 1 μg/mL for 20 minutes at 37°C on a shaker to stain nuclei. Slices were carefully mounted with a proprietary solution, DeepLabel XClarity, and allowed to set overnight before imaging.

The inhibitory synaptic stain was performed in the same way, with a different antibody combination. Primary antibodies were anti-VGAT (chicken, Synaptic Systems 131 006) and anti-Gephyrin (mouse, Synaptic Systems 147 111). Secondary antibodies consisted of goat anti-chicken Alexa Fluor Plus 647 (Invitrogen A32933), and goat anti-mouse Alexa Fluor Plus 488 (Invitrogen A32723) with DAPI included as above. 1:2000 dilution for these antibodies also obtained good staining.

Imaging of these samples was performed on the Zeiss AiryScan confocal microscope system (Carl Zeiss, Jena Germany). We took z-stacks with frames 0.2 μm apart at 63x magnification, comprising 16 such frames for a total thickness of 3 μm. With X/Y dimensions of 72 x 72 μm, the total volume surveyed for these experiments was 15,552 μm3.

### Synaptic imaging analysis

Raw images were passed through the AiryScan image processing system native to Zeiss’s Zen Black, increasing their resolution by roughly twofold. Using FIJI (Schindelin et al., 2012) and MATLAB, these images were then converted into single-channel .TIFF files, scaled to normalize for intensity over depth, and deconvoluted with a point-spread-function inferred from TetraSpeck 100 nm fluorescent microspheres (Thermo Fisher Scientific T7279, Waltham MA) imaged with the same optical setup and processing. To identify synaptic puncta in our channels of interest, then, we used a system of thresholding based on an estimate of the likely maximal brightness of noise in our image. A skewed normal distribution was fitted to the histogram of pixel intensity values and the percent likelihood that a given intensity value belonged to this distribution was computed. The inverse of this, the likelihood that a value was part of the signal, was transformed into an intensity threshold for peak detection. An experimenter blinded to condition settled on a threshold for each stack that excluded objects that were clearly out of focus. Any object or part of an object that was dimmer than that was not considered a potential region of interest (ROI).

Thresholded ROIs had edges defined by a second, slightly lower, threshold (Wise, Escobedo-Lozoya et al., 2023). They were then further expanded by three pixels in each direction to make certain that we were not missing pixels that were very close to the threshold. ROIs were resegmented using a watershed algorithm into component puncta objects, then restricted to the pixels which were within 30% of the difference between the puncta’s peak and the local background. ROIs that were thereafter under 8 pixels (0.013 μm^3^) in volume were excluded from the final dataset. Finally, the colocalization of each punctum with its synaptic partner was assessed. Puncta were considered colocalized if they were within 2 pixels of a potential partner. These colocalized puncta are the population reported here for their properties - density and volume.

Cell counts were achieved with 10x imaging of layer 5 of somatosensory cortex and surrounds, where we made all our measurements. A 461 x 461 μm area was imaged for its full extent in z for each sample, and nuclei were classified and counted in CellProfiler (Stirling et al., 2021). Density was then calculated for each stack, one per imaged slice from the synapse imaging dataset.

## RESULTS

### Calcium transients demonstrate a progressive loss of activity homeostasis with prolonged deprivation

Spontaneous activity in cortical organotypic slice cultures becomes progressively more complex and frequent over the first few weeks *in vitro* (Johnson and Buonomano, 2007), characterized by periodic population firing events, termed ‘up states,’ in which neurons depolarize to make activity propagation more likely (Cossart et al., 2003). These up states were evident in GCaMP6f fluorescence recordings of network activity, monitored at 15-18 days in vitro (Equivalent Postnatal Day; EP 22-25; **Figure 1A & 1B**, left). Four different measures were used to quantify network activity: synchrony was quantified as between-cells correlation at zero lag between cell pairs within the period of each up state, then averaged across all cells and all up states; stereotypy was the within-cell correlation of a cell’s activity in one up state with its activity in other up states, calculated per cell on each pair of up states then averaged across all up state pairs and all cells; up state frequency was simply the count of up states within an average minute; up state duration was assessed as the time between the first period of increasing slope in an up state and the last. Activity in healthy slices was characterized by several consistent features: 1) a modest frequency of network up states (mean ∼1.8 up states per minute) remaining relatively constant from EP22 and EP25; 2) calcium transients in simultaneously imaged cells, though coarsely synchronous in that they were grouped into 5-30s long up states (mean 8.7s), were not tightly synchronized at more rapid time scales (synchrony measures below 0.6, **Figure 1E**); and 3) activity in simultaneously recorded cells often occurred in a different order in different up states (low levels of stereotypy, **Figure 1F**), with many cells active in some up states and not others. This suggests a circuit that has achieved a relatively balanced activity state in culture following the regrowth of severed connections from the dissection process.

**Figure 1:**
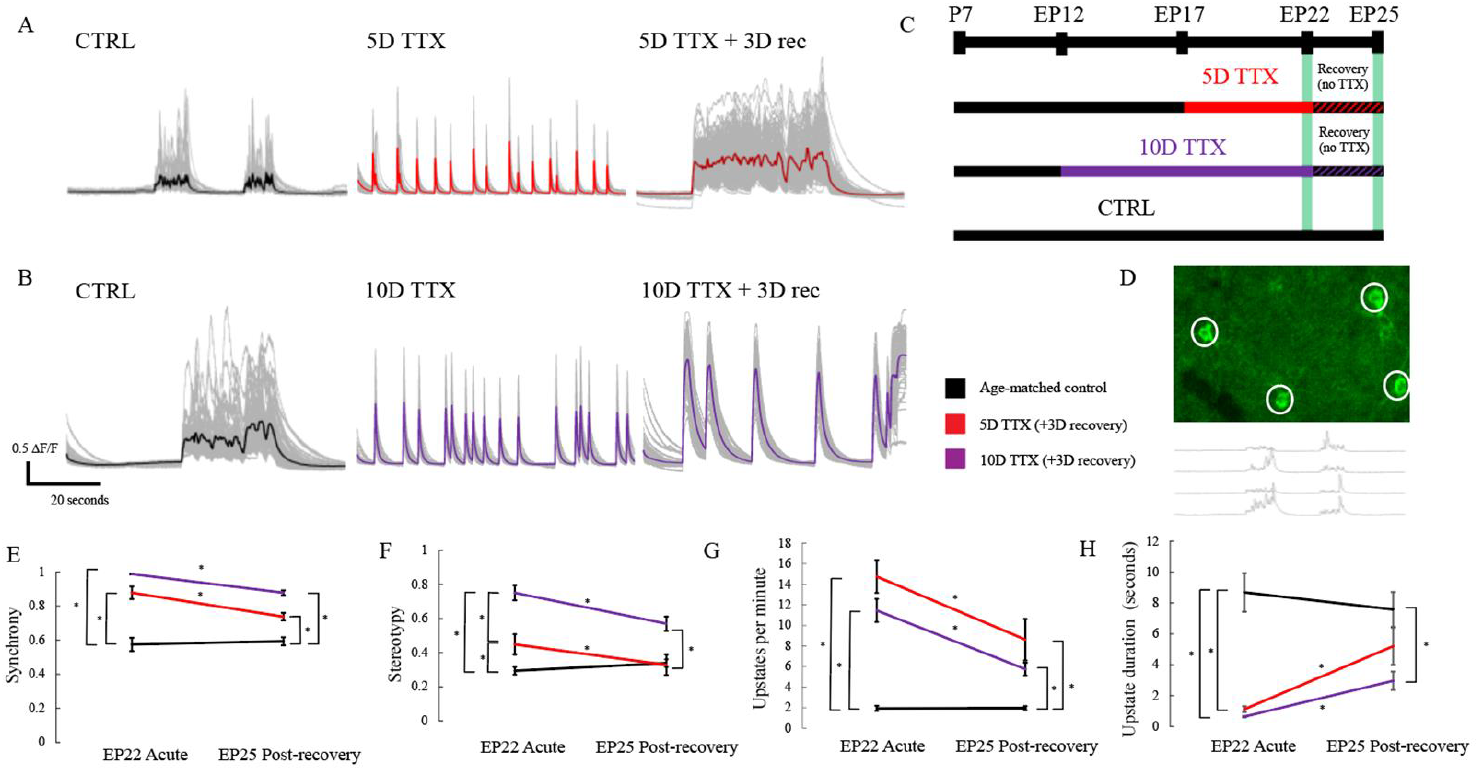
Population activity in organotypic cortical slice cultures is hyperactive and hypersynchronous upon release from prolonged activity deprivation and following 3 days recovery. **A**. Traces (mean in color, individual cells in gray) show calcium transients recorded from individual cells after normal activity (CTRL, black, left), after release from 5D of TTX (center, red), or after recovery for 3D in normal media before imaging (right, red). **B**. Same as A but following 10D of TTX exposure (purple). **C**. Experimental timeline (n = 23 EP22 CTRL, 27 EP25 CTRL, 16 EP22 5D TTX, 19 EP25 TTX 3D recovery, 14 EP22 10D TTX, and 12 EP25 10D TTX 3D recovery slices). **D**. Representative single frame CTRL calcium imaging with white circles representing cellular ROIs with high GCAMP6f fluorescence traces below show these four cells over 1 minute of imaging. **E**. Synchrony, averaged across all pairs and up states. Black: age-matched control; red: 5D of TTX; purple: 10D of TTX, immediately after activity deprivation (EP22 acute) and following an additional 3D of recovery in normal media (EP25 post-recovery). Stars: p < 0.05 (2-way ANOVA, Tukey’s post hoc test). **F**. Stereotypy, calculated as autocorrelation at zero lag of each cell with its activity across up states, averaged across all cells. **G**. Frequency of activity, as up states per minute. **H**. Average up state duration, across all up states in the 10-minute recording window, in seconds.

Following prolonged activity blockade, organotypic cultures exhibited a highly stereotyped and synchronized state of epileptiform activity (**Figure 1A & 1B**, middle) as previously described for slice culture and dissociated cultures (Koch et al., 2010; Ramakers et al., 1990; Trasande and Ramirez, 2007). Up states were observed significantly more frequently after silencing, shifting from a mean of 1.9 events/minute to 14.7 events/minute at 5D TTX to 11.5 events/minute with 10D TTX. When one neuron fired, most or all other neighbors rapidly became active as well, leading to a synchrony approaching a perfect correlation of 1 (mean 0.88 5D TTX, 0.99 10D TTX). Each of the up states involved a highly stereotyped burst, where each up state appeared as a sudden, brief flash (stereotypy mean 0.45 5D TTX, 0.75 10D TTX; an increase of 55% and 158% respective relative to control activity). The up state duration dropped from mean 8.7s in control activity to 1.1s following 5D TTX and to 0.6s following 10D TTX. This highly synchronous rebound firing after activity deprivation occurred in every slice examined, likely reflecting persisting homeostatic drive accumulating in the absence of feedback during tetrodotoxin silencing. The greater increase in synchrony and stereotypy after 10D TTX suggests that homeostatic changes continue to occur even after the initial 5D of deprivation.

One exception to this pattern was that up state frequency was increased more following 5D of TTX than 10D of TTX, but this difference was not significant (5D *vs*. 10D TTX acute p = 0.26 Tukey’s post hoc test, 5D *vs*. 10D TTX). The duration of up states was also not significantly different immediately after 5 and 10D TTX (p = 0.99). In both cases, post TTX activity occurs in short synchronous bursts. If the 10D TTX bursts had higher overall firing rates, however, this might lead to a longer period of inactivity between bursts. The calcium imaging approach used had neither the temporal resolution nor the precise calibration to be able to directly infer action potential firing rates from calcium transients.

The greater hyperexcitability seen after 10D TTX might also be associated with a slower or less complete recovery following the period of deprivation. To assess this, we monitored activity following 3 days in normal ACSF. Each of the four metrics examined showed some degree of recovery, but most did not recover fully after 3 days and in general recovery was more complete following 5D TTX than following 10D TTX. For example, after 5D TTX, synchrony recovered to an average synchrony of 0.74, approximately halfway from the acute value of 0.88 to the control value of 0.59, but this remained significantly different from control (p = 0.003 Tukey’s post hoc test). While after 10D TTX, synchrony recovered to an average of 0.88, which was only about a quarter of the way from the acute value of 0.99 to the baseline value. This synchrony after recovery was significantly different from baseline (p < 0.01) as well as from the recovery after 5D TTX (p = 0.01). This differential recovery was even more striking for stereotypy which recovered completely following 5D TTX (mean 0.33 *vs*. 0.34 CTRL, p = 0.99), but only partially following 10D TTX (mean 0.57 *vs*. 0.34 CTRL, p < 0.01). Up state duration also showed more complete recovery following 5D TTX than 10D TTX (CTRL mean 7.6s, 5D TTX rec 5.2s and 10D TTX rec 3.0s). The 5D TTX recovery was not significantly different from Control (p = 0.43), but the 10D recovery was still significantly different from CTRL (p = 0.01). As for the acute effects, the exception to this pattern was the up state frequency where both after 5D TTX and after 10D TTX the recovery was incomplete (p < 0.01 and p = 0.03 *vs*. CTRL respectively).

Qualitatively, up states after recovery from 5D TTX sometimes resembled the control state (Fig 1A), although this was not always the case. In contrast, 3 D recovery was usually insufficient after 10 D TTX to restore a more asynchronous pattern of network activity.

Overall, calcium imaging reveals that populations of neurons transiently overshoot their homeostatic set point when deprived of activity for a prolonged period and this hyperexcitability is more extreme following a longer period of deprivation. A three-day period of recovery in normal media results in an incomplete recovery that is correspondingly less restorative when the deprivation is more prolonged.

### Longer periods of recovery produce a more complete return to baseline activity

Because 3D was clearly insufficient, we investigated whether longer recovery could better ameliorate the lasting impact of silencing (**Figure 2**). Specifically, we extended the recovery period after 10D of TTX to 7 or 10D. For synchrony (P32 CTRL *vs*. 10D TTX 10D recovery, p = 0.56), stereotypy (p = 0.37), and up state frequency (p = 0.99), there was complete recovery back down to age-matched control values. Critically, our results do not imply that recovered circuits are in all ways identical to their original state, but only that our population activity metrics capturing average events grossly recover to baseline level. A circuit of this type could arrive at exactly the same overall balance of activity but still be more fragile to additional insult or strong stimulus.

**Figure 2:**
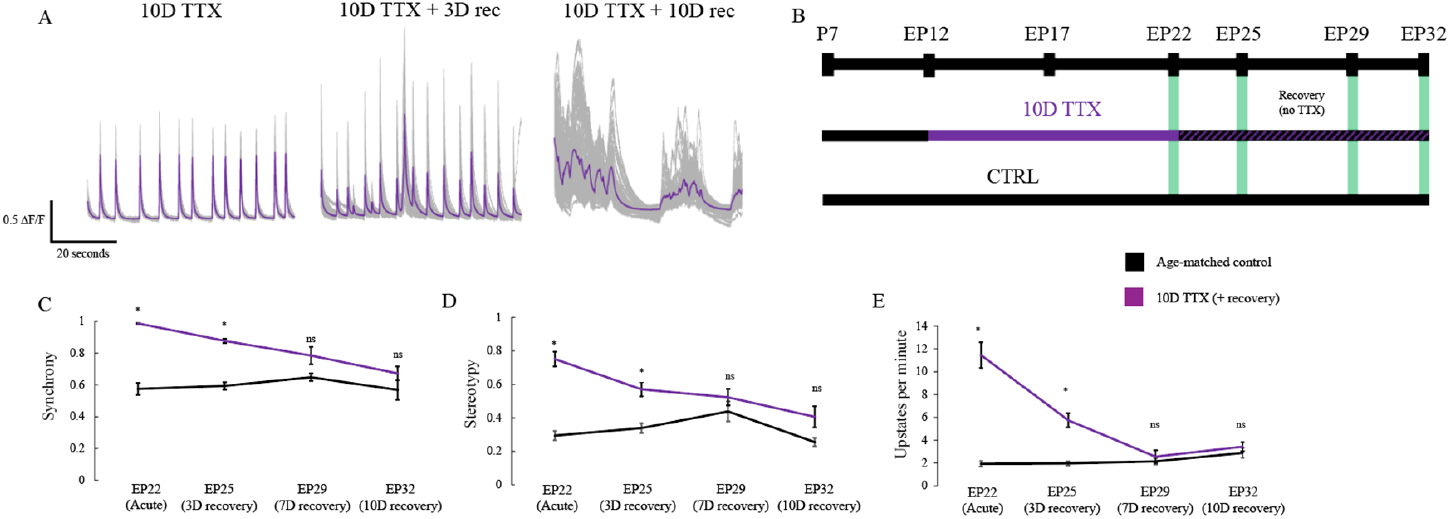
Longer recovery following prolonged deprivation does eventually equilibrate in healthy activity patterns. **A**. Traces (mean in purple, individual cells in gray) show calcium transients recorded from individual cells after normal activity (10D TTX, left; same group as in Fig 1), after 3D of recovery in normal media following 10D of TTX (center, as in Figure 1), after 7D recovery (n = 7 for EP29 age-matched CTRL slices, 10 10D TTX 7D recovery, not shown), and after recovering for 10D in total (right, n = 10 for EP32 age-matched CTRL slices, 16 10D TTX 10D recovery). **B**. Experimental timeline. **C**. Synchrony progressively recovers by 7D, fully recovered by 10D. Stars: p <\ 0.05 (2-way ANOVA, Tukey’s post hoc test). **D**. Stereotypy, otherwise as in **C. E**. Up state frequency per minute, otherwise as in **C**.

### Hyperexcitability is most pronounced when activity blockade is initiated prior to the end of the second postnatal week

Homeostatic plasticity is regulated developmentally, with stronger plasticity observed earlier in development (Desai et al., 2002). The maladaptive response to activity deprivation in vivo is also strongly regulated by development and does not lead to recurring seizures if initiated after the end of the second postnatal week of development (Galvan et al., 2000; Lee et al., 2008). To see if this correlated with the developmental sensitivity to epileptiform activity in slice culture, we initiated 5 days of activity deprivation beginning at progressively later equivalent ages from EP12 to EP22 and measured the resulting hyperactivity at EP17, EP22 and EP27 (**Figure 3**). Synchrony (**Figure 3C**) and stereotypy (**Figure 3D**) indices revealed greater hyperactivity for deprivation begun earlier. Synchrony had significantly less effect by P27 with 5D TTX when compared with P17 (p < 0.01 Tukey’s posthoc test), whereas stereotypy was actually no longer statistically different from age-matched control by P22 TTX (from p < 0.01 at P17 to p = 0.99 at P22 and p = 0.78 at P27). For peak frequency, the effect of development was non-monotonic, with the largest effect evident at the intermediate age (**Figure 3E**). This may reflect an interaction between progressively enhanced activity which increases the number of firing events and progressively lengthened intervals between high activity bursts, which tends to reduce this measure.

**Figure 3:**
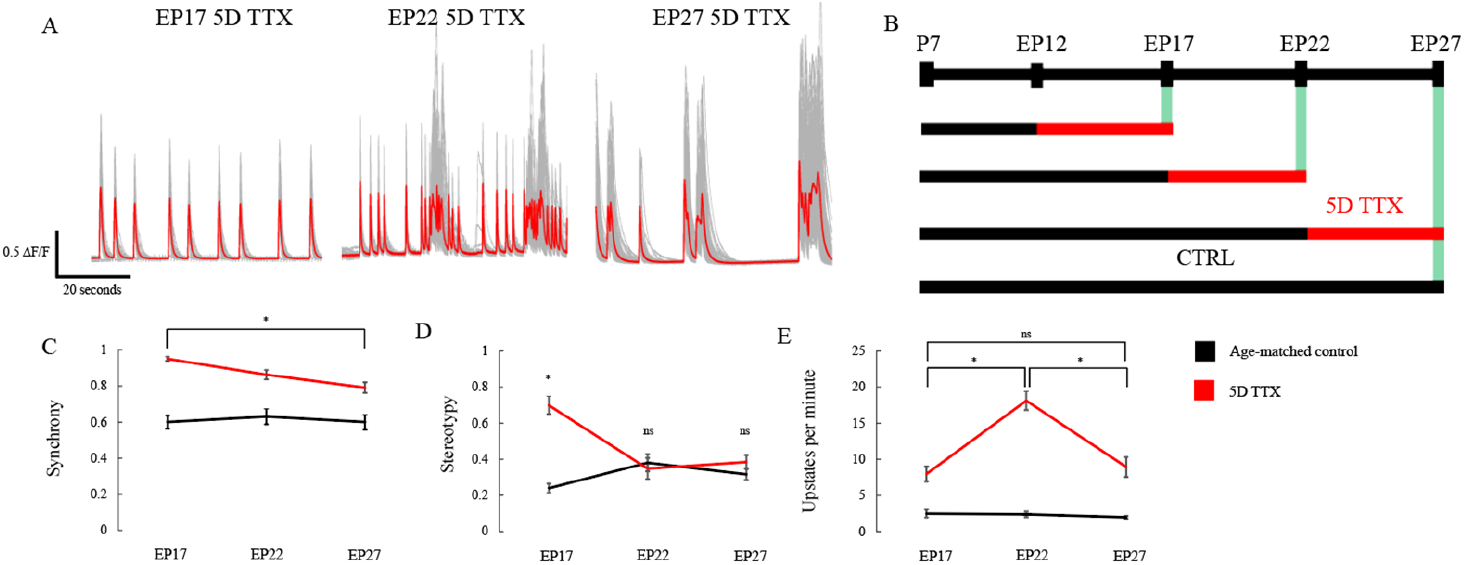
The overshoot following 5D of TTX is age-dependent, with the strongest effects occurring when initiated prior to the end of the second postnatal week. **A**. Traces (mean in color, individual cells in gray) show calcium transients recorded from individual cells after 5D of TTX (ending EP17, left, n = 15 CTRL and 18 5D TTX slices; EP22, middle, n = 15 CTRL and 16 5D TTX slices; EP27, late, n = 23 CTRL and 26 5D TTX slices). **B**. Experimental timeline. **C**. Synchrony overshoot is strongest at EP17 and is significantly reduced by EP27. Stars: p < 0.05 (2-way ANOVA, Tukey’s post hoc test). **D**. Stereotypy recovers back to baseline conditions by EP22. **E**. Up state frequency displays a non-monotonic relationship.

### Pharmacological block of NMDA receptors suggests Hebbian mechanisms make little contribution to circuit hyperexcitability

Synaptic scaling at neocortical synapses is known not to require activation of NMDA receptors because blockade of NMDA receptors does not itself produce scaling of AMPA-mediated transmission (Turrigiano et al. 1998). However, following release from activity blockade, elevated network activity might lead to enhanced NMDA-dependent Hebbian plasticity, thus “locking in” the initial homeostatic increase in synaptic strength. To test for this possibility, we compared activity in normal ACSF following either three days of recovery with NMDA receptors blocked with 2-amino-5-phosphonopentanoic acid (APV), or with NMDA receptors available for normal activation. The concentration of APV used (50 μM) is known to be sufficient to block long-term potentiation (LTP) at neocortical synapses (Crair & Malenka, 1995; Sjöström et al., 2001). We observed comparable recovery across each of our three metrics regardless of whether NMDA receptors were blocked or not (**Figure 4**, 10D TTX 3D APV *vs*. 10D TTX 3D recovery synchrony p = 0.99, stereotypy p = 0.98, up state frequency p = 0.87). In both cases, slices failed to recover back to age-matched baseline (10D TTX 3D APV *vs*. P25 CTRL synchrony p < 0.01, stereotypy p = 0.04, up state frequency p = 0.04). Since recovery was not enhanced by blocking NMDA receptors, we conclude that continued network hyperactivity after 3D of recovery is a persistent consequence of homeostatic plasticity, rather than a consequence of Hebbian plasticity engaged during the subsequent period of highly synchronous activity.

**Figure 4:**
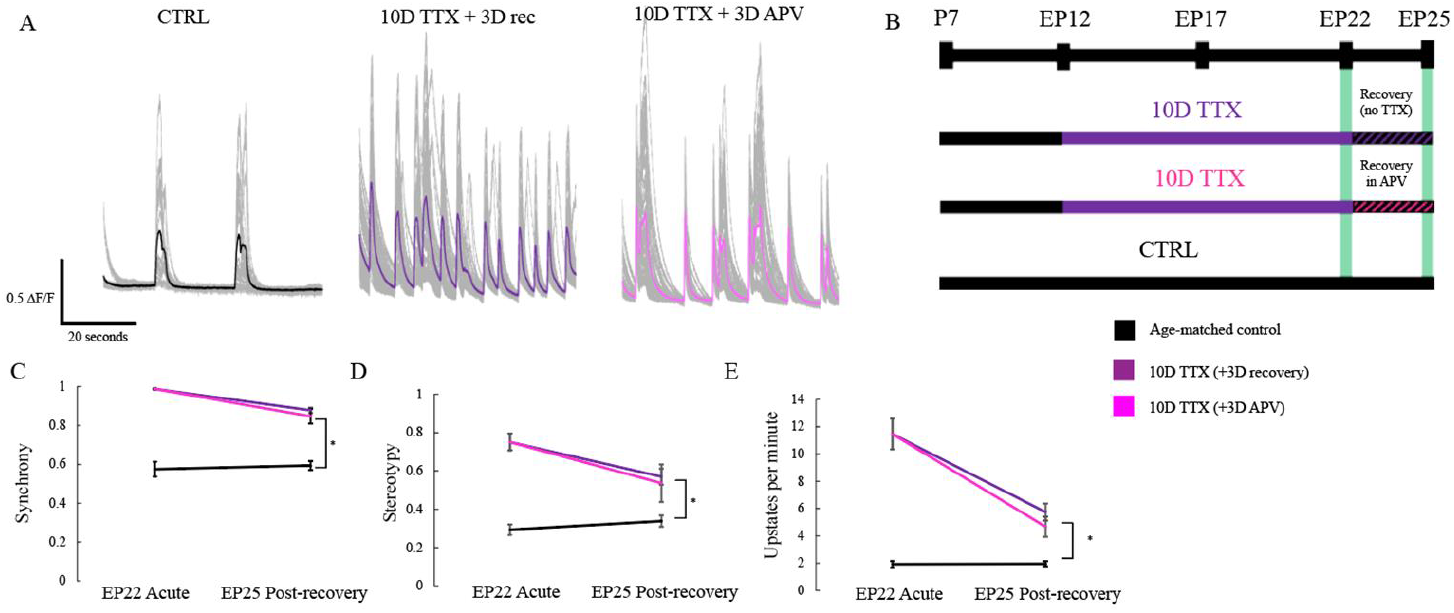
Maladaptive plasticity does not require activation of NMDA receptors during the recovery period. **A**. Traces (mean in color, individual cells in gray) show calcium transients recorded from individual cells after normal activity (CTRL, black, left; samples from Figure 1), after 3D of recovery in normal media following 10D of TTX (center, purple, from Figure 1), or after recovering in 3D APV before imaging (right, pink; n = 10 10D TTX 3D APV slices). **B**. Experimental timeline. **C**. Synchrony remains significantly elevated after 3D but this is not significantly altered by the presence or absence of APV during the recovery period. Stars: p < 0.05 (2-way ANOVA, Tukey’s post hoc test). **D**. Stereotypy, otherwise as in **C. E**. Up state frequency per minute, otherwise as in **C**.

### Intrinsic excitability of L5 pyramidal neurons is slightly increased, and input resistance is increased two-fold, with prolonged deprivation

In order to begin to reveal which cellular and synaptic mechanisms of homeostatic plasticity might contribute to persistent hyperexcitability, we examined intrinsic excitability of pyramidal neurons after a 3D recovery period following TTX silencing for 5 or 10 days. Specifically, we measured frequency-current (FI) curves over a range of 0 pA to 450 pA during whole cell current clamp recordings (**Figure 5**). Many of our 10D TTX 3D recovery cells inactivated after a few action potentials at higher current steps, so we relied instead on instantaneous firing rate between the second and third spike (FI ⅔, **Figure 5B**). Other measures of firing we calculated included the interval between the first and second spike, and the average number of action potentials (excluding periods of inactivating firing), yielded similar conclusions but were noisier (not shown; FI 1/2 and standard FI). The instantaneous FI from spike 2-3 showed a main effect for group and for current level, as well as a significant interaction effect of group and current levels (2-way repeated measures ANOVA, F = 3.0, p = 0.04 for between-subject effect of condition, F = 173.3, p < 0.01 for within-subject effect of current step, F = 2.8, p = 0.02 for interaction of group and current). Post-hoc tests reveal an increase in firing from CTRL to 10D TTX 3D recovery at 300, 350, 400, and 450 pA (Tukey’s; p = 0.03, p = 0.01, p = 0.01, p = 0.02 respectively), though 5D TTX was never significantly different than CTRL (closest was p = 0.10 at 400 pA). We made some attempt to quantify FI behavior in neurons acutely withdrawn from TTX, but 10D of TTX consistently produced cells that inactivated even after a single spike, making quantitative comparisons difficult.

**Figure 5:**
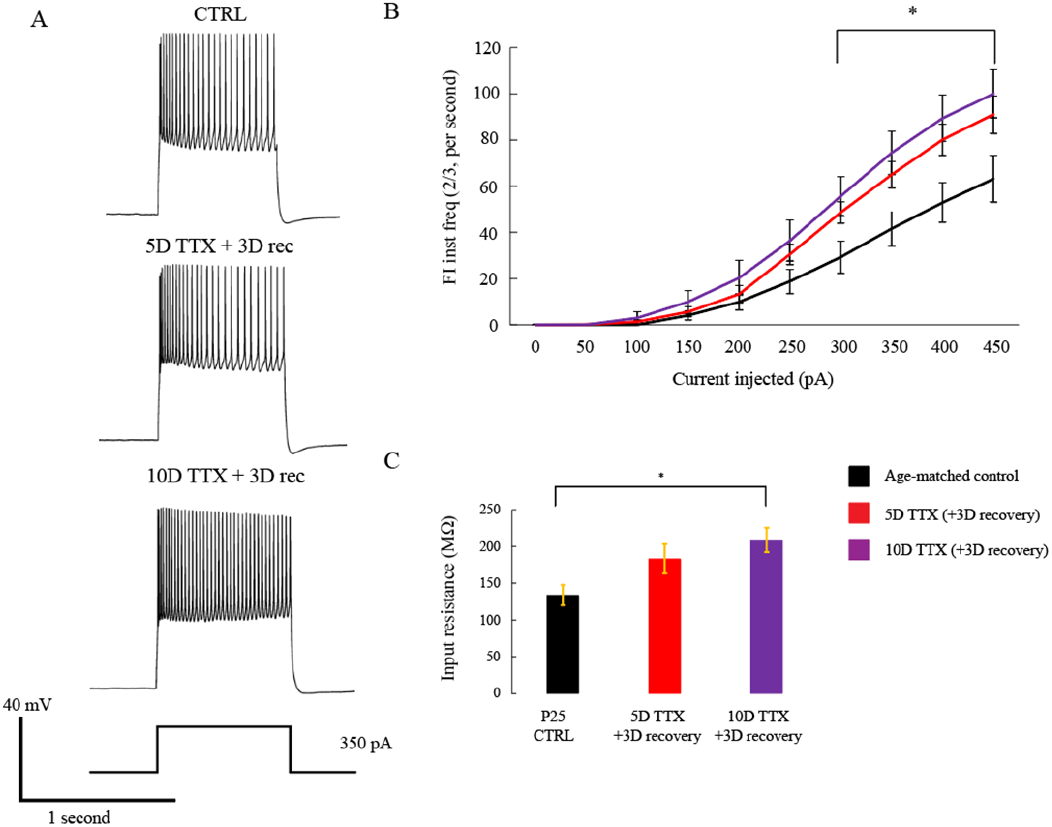
Intrinsic excitability measurements from L5 pyramidal neurons reveal an increase in instantaneous firing rate and a large increase in input resistance, with no persistent change in rheobase. **A**. Single trial traces of current-clamped neurons in each condition most closely approximating the mean firing observed in each condition (EP25 age-matched control, n = 14 cells; EP25 5D TTX 3D recovery, n = 12 cells; and EP25 10D TTX 3D recovery, n = 15 cells) with 350 pA stimulation. **B**. Instantaneous firing rate over current injection steps from 0 pA to 450 pA. **C**. Input resistance from recorded cells, averaged across trials. Star represents significance following 1-way ANOVA and Tukey’s honestly significant difference test of p < 0.05.

In addition to greater suprathreshold firing, deprivation made neurons more excitable by increasing input resistance at rest. There was a progressive increase in input resistance with increasing deprivation duration (**Figure 5C**, 1-way ANOVA, F = 5.23, p = 0.01) that roughly doubled and was significantly different from age-matched control by 10D of TTX (CTRL mean 133.8 MΩ, 5D TTX 3D recovery 183.2, 10D TTX 3D recovery 208.9; p = 0.13 for CTRL *vs*. 5D TTX 3D recovery, p = 0.01 for CTRL *vs*. 10D TTX 3D recovery). Increased input resistance leads to greater firing because a given level of current injection produces stronger depolarization. We calculated the extrapolated intercept of the FI curve as the rheobase, the threshold current for a single spike, and found it to not significantly differ across conditions (1-way ANOVA, F = 0.75, p = 0.48). It may be that a small difference, less than 50 pA, does exist but that our FI step size was too crude to reveal it. It appears that the increase in input resistance may have contributed to the increase in instantaneous firing we see.

Even given 3D of recovery it is clear that intrinsic excitability in our cultures did not fully recover. Unlike in our calcium metrics, 5D TTX displayed a similar shift in F/I dynamics to 10D TTX. However, input resistance shows a progressive change with increasing silencing duration.

### A decrease in excitatory synapse density, but larger decrease in inhibitory synapse density, indicate a shift in E/I balance

In order to measure the impact of progressive deprivation and recovery on cortical synapses, we visualized synapse density and size using super-resolution microscopy (Zeiss Airyscan, see Methods). Putative excitatory synapses were defined by the colocalization of presynaptic puncta (Bassoon) with postsynaptic puncta (excitatory-specific PSD-95). This method has recently been calibrated and compared to electron microscopy in an analogous preparation (Wise, Escobedo-Lozoya et al., 2023).

Most prior studies of deprivation-induced changes at cortical synapses have emphasized the role of increased excitatory postsynaptic strength (O’Brien et al., 1998; Turrigiano et al., 1998). Our measure of postsynaptic size (2D cross-sectional area of colocalized PSD-95) revealed a subtle change. Size exhibited a progressive increase as slices were silenced with TTX (**Figure 6B**, a non-significant 15% increase from CTRL to 10D TTX 3D recovery) (1-way ANOVA: F = 1.41, p = 0.26). The divergence between this result and consistent reports of increased postsynaptic size after acute deprivation, including in a recent study also looking at prolonged TTX in cortical organotypic culture (Wise, Escobedo-Lozoya et al., 2023), likely reflect some degree of reversibility of synaptic size during a period of recovery.

**Figure 6:**
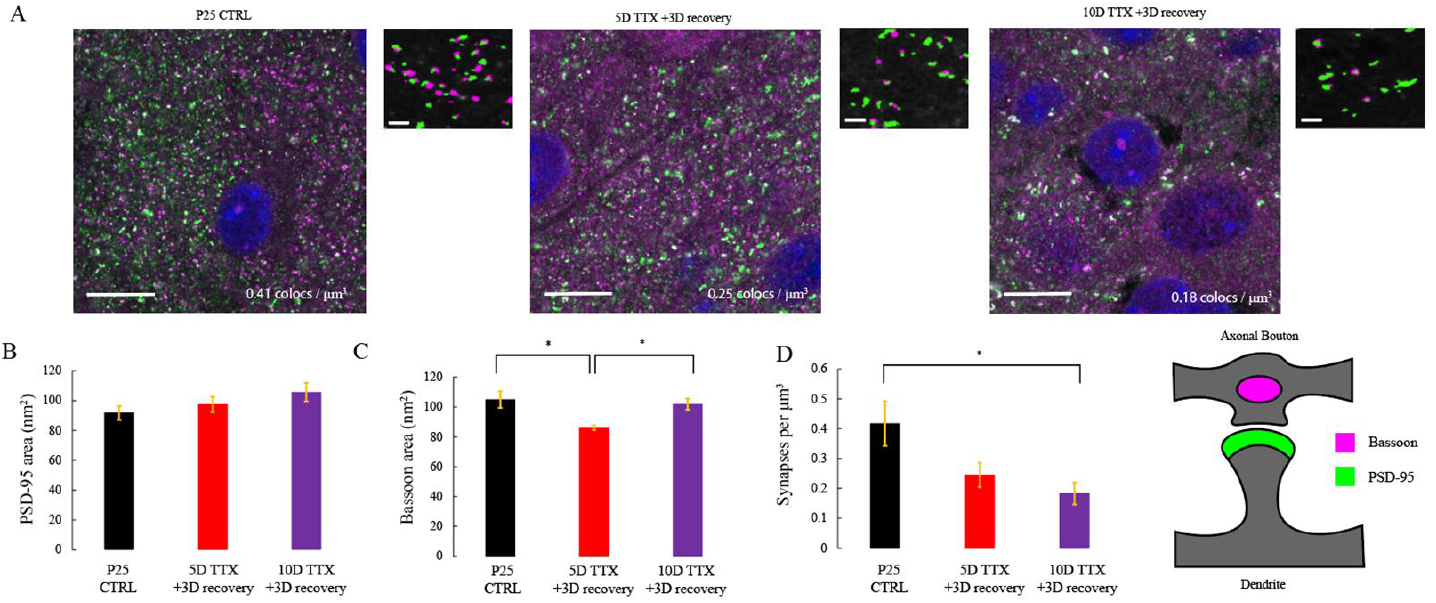
Excitatory synapse number progressively declines after 10D of TTX activity deprivation. **A**. Representative example images, each a single z-frame, of organotypic slice culture tissue stained with immunofluorescent markers for the postsynapse (PSD-95, green), and presynapse (Bassoon, magenta), schematized on the far right. Scale bar on large image is 10 μm. Right panel of each duo shows a zoomed in (100 pixel x 100 pixel, or 72 μm square) view of a single area with thresholded colocalized puncta pseudo colored, and background of the Bassoon stain in greyscale. Scale bar on small image is 1 μm. n = 7 EP25 CTRL slices, 11 EP25 5D TTX 3D recovery, 8 EP2510D TTX 3D recovery. **B**. Average 2D cross-sectional area (in frame with brightest pixel) of PSD-95 puncta colocalized with synaptic partners within a 16.59 mm^3^ area. Compared are EP25 age-matched untreated slices with 5D TTX + 3D recovery and 10D TTX +3D recovery slices. CTRL mean 91.8 nm^2^ at CTRL, 5D TTX 3D recovery 97.5 nm^2^, 10D TTX 3D recovery 105.7 nm^2^. **C**. Average 2D area of Bassoon puncta colocalized with synaptic partners. CTRL mean 105.1 nm^2^ at CTRL, 5D TTX 3D recovery 86.3 nm^2^, 10D TTX 3D recovery 101.9 nm^2^. **D**. Presumptive synapse density, estimated as a count of PSD-95 puncta that are colocalized with Bassoon partners, across conditions. CTRL mean 0.41 colocalizations/μm^3^, 5D TTX 3D recovery 0.25 CLs/μm^3^, 10D TTX 3D recovery 0.18 CLs/μm^3^.

The relationship between the progressive silencing and the size of the presynaptic Bassoon puncta was more complex and non-monotonic. Bassoon 2D area first decreased after 5D of TTX but had returned to control levels following 10D of treatment and 3D of recovery, resulting in a significant modification of size with treatment (**Figure 6C**, F = 5.95, p < 0.01). Post hoc tests of condition differences were significant for the 5D TTX condition relative to CTRL (p = 0.01) and to 10D TTX (p = 0.04), but CTRL and 10D TTX were not significantly different (p = 0.87). This does not accord with the results of deprivation prior to recovery, which have found matching increases in both pre- and postsynaptic size (Mitra et al., 2011; (Wise, Escobedo-Lozoya et al., 2023). This discrepancy could reflect the effects of elevated activity during the recovery period.

Presumptive excitatory synapses were also reduced in density over increasing TTX duration (**Figure 6C**). Synapse density, defined as the number of PSD puncta colocalized with a Bassoon presynaptic partner per unit area, dropped sharply with increasing duration of silencing. This 57% loss of excitatory synapses was significant by 10D of TTX, whereas a 42% decrease was not significant at 5D of TTX (1-way ANOVA with THSD post hoc, F = 5.10, p = 0.01; p = 0.06 for CTRL *vs*. 5D TTX 3D recovery, p = 0.01 for CTRL *vs*. 10D TTX 3D recovery, p = 0.64 for 5D TTX 3D recovery *vs*. 10D TTX 3D recovery). A comparable decrease was recently observed acutely after 5D deprivation without recovery (Wise, Escobedo-Lozoya et al., 2023). This may suggest that the network is unable to restore excitatory synapse density within three days.

Network activity depends not only on intrinsic excitability and excitatory synaptic strength and number, but also critically on the strength and number of inhibitory synapses. Inhibitory synapse density was previously shown to decrease with silencing (Chattopadhyaya et al., 2004), with concomitant synaptic weakening (Kilman et al., 2002). We labeled inhibitory presynaptic terminals with VGAT and postsynapses with gephyrin. Previously observed reductions in size were not observed following deprivation with a period of recovery. Here, postsynaptic gephyrin puncta were not significantly altered in size (**Figure 7B**, 1-way ANOVA F = 0.62, p = 0.54), while presynaptic VGAT puncta were larger following 10D silencing (**Figure 7C**, 1-way ANOVA: F = 7.42, p < 0.01; p = 0.71 for CTRL *vs*. 5D TTX 3D recovery, p = 0.02 for CTRL *vs*. 10D TTX 3D recovery, p < 0.01 for 5D TTX 3D recovery *vs*. 10D TTX 3D recovery). These modest changes in inhibitory synapse size were dwarfed by a profound and progressive reduction in inhibitory synapse number (**Figure 7D**, 67% decrease from CTRL to 10D TTX, 51% decrease at 5D TTX), which was highly significant as a 1-way ANOVA (F = 8.94, p < 0.01) and represented a significant shift with deprivation from age-matched CTRL (p = 0.02 *vs*. 5D TTX 3D recovery; p < 0.01 *vs*. 10D TTX 3D recovery) though not between the two deprived conditions (p = 0.53). These results indicate that the loss of inhibitory synapses is greater than the loss of excitatory synapses, suggesting a reduction in the E/I balance likely to favor the emergence of epileptiform activity and circuit hyperexcitability (Nelson and Valakh, 2015).

**Figure 7:**
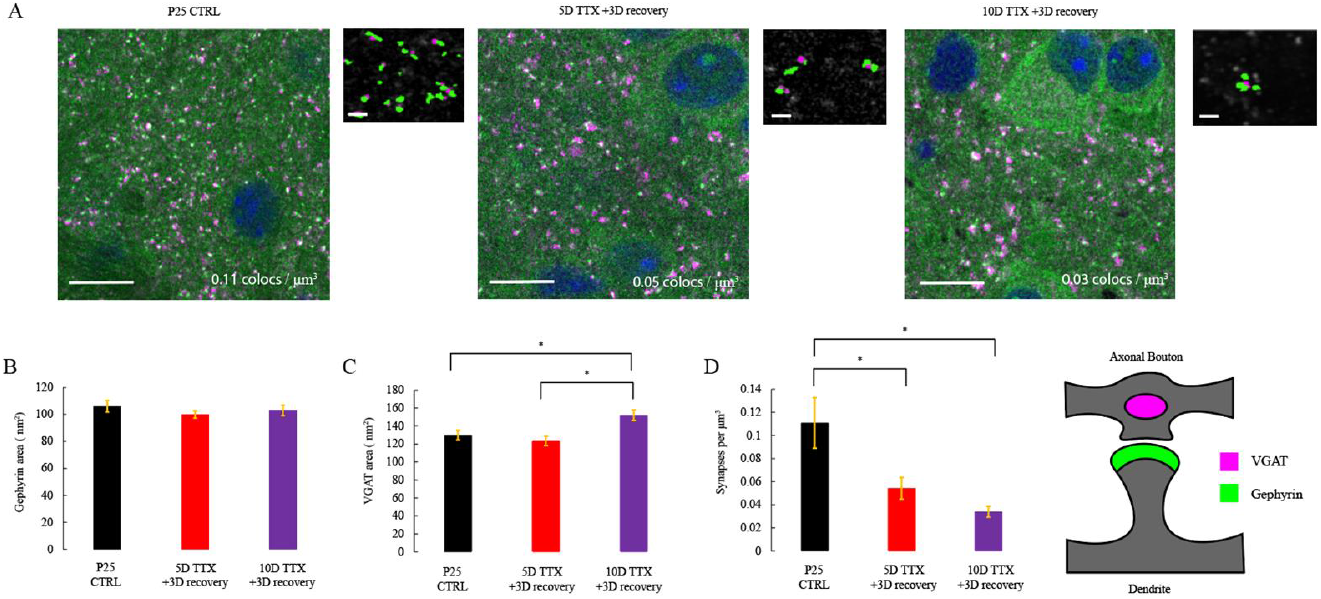
Inhibitory synapse density progressively declines with increasing duration of silencing as presynaptic inhibitory protein marker increases in size. **A**. Representative z-frames of slices, as in Figure 4, stained at the postsynapse (gephyrin, green), and presynapse (VGAT, magenta). Scale bar on large image is 10 μm. Right panel of each duo shows a zoomed in (100 pixel x 100 pixel, or 72 μm square) view of a single area with thresholded colocalized puncta pseudo colored, and background of the gephyrin stain in greyscale. Scale bar on small image is 1 μm. n = 12 EP25 CTRL slices, 12 EP25 5D TTX 3D recovery, 14 EP25 10D TTX 3D recovery. **B**. Average 2D cross-sectional area (in frame with brightest pixel) within a 0.72 μm x 0.72 μm x 3 μm stack of gephyrin puncta colocalized with synaptic partner, as in Figure 5 (age-matched control, 5D TTX + 3D recovery and 10D TTX +3D recovery). CTRL mean 105.9 nm^2^, 5D TTX 3D recovery 99.9 nm^2^, 10D TTX 3D recovery 103.0 nm^2^. **C**. Average 2D area of VGAT puncta colocalized with synaptic partner. CTRL mean 129.9 nm^2^, 5D TTX 3D recovery 123.5 nm^2^, 10D TTX 3D recovery 151.9 nm^2^. **D**. Gephyrin colocalized puncta density. CTRL mean 0.11 colocalizations/μm^3^, 5D TTX 3D recovery 0.05 CLs/μm^3^, 10D TTX 3D recovery 0.03 CLs/μm^3^.

### No cell death induced by treatment with tetrodotoxin

Organotypic slices cultured in the presence of TTX exhibited an obvious visual thinning of the tissue that, if shown to indicate significant cell loss, might account for some or all of the decrease in synapse count (Dingledine et al., 2014). We counted nuclei stained with a DAPI stain from layer five of somatosensory cortex at 10x on a confocal microscope (Supplemental **Figure 1**), over a z-stack containing the whole thickness of the same slices used for synapse imaging, on the three conditions of increasing silencing duration assayed prior. There was no difference across conditions in cell count (1-way ANOVA, F = 0.61, p = 0.55). This measure offers no insight into whether there is a change in the relative mixture of different cell types (such as a loss of neurons accompanied by an increase in astrocyte density) because DAPI binds to DNA in the nucleus of all cells. Nevertheless, we can be confident that massive cell death did not occur during our deprivation experiments.

## DISCUSSION

Prior studies of the response of central circuits to activity deprivation have mostly focused on brief, rapidly reversible homeostatic changes that normally allow circuits to maintain stable firing rates. Here, we have focused instead on the progressive and slowly reversing compensatory plasticity that eventually leads to hypersynchronous seizure-like activity. The observed plasticity shares many features in common with the seizure disorder that develops *in vivo* during chronic activity blockade (Galvan et al., 2000; Lee et al., 2008). First, as *in vivo*, the observed network activity became progressively stronger during longer periods of deprivation. Second, as in vivo, the effect of deprivation was strongest when begun prior to the end of the second postnatal week, a period corresponding to a period of heightened (Aaberg et al., 2017; Silverstein and Jensen, 2007) risk of developing seizures in human patients. Third, the activity is slow to recover, showing incomplete recovery after 3 days and only showing full recovery after a period equal to that of the deprivation (10 days). Limitations of the ability to maintain healthy slice cultures for very long periods limited our ability to look at longer periods of deprivation and longer periods of recovery. *In vivo*, seizures take 10-14 days of disruption to develop but then do not recover. However, this may involve many brain regions and perhaps require changes set in motion once seizures begin. Although we could not rule out such lasting effects of seizures once begun *in vivo*, we did rule out the idea that persistent hyperactivity was due to Hebbian plasticity induced by epileptiform rebound bursting. Blocking NMDA receptors during this period did not lead to more rapid recovery, arguing against the idea that epileptiform activity was “locked in” by Hebbian LTP during synchronous network activity.

### Activity deprivation, homeostatic mechanisms, and seizure disorders

The effects of activity deprivation may have broader application to more common forms of seizure disorders. Epilepsy can begin following stroke or injury (Curia et al., 2016), which can trigger additional waves of cell death in cortex, especially in young animals (Bittigau et al., 2003). Much is still unknown about how seizures result from trauma. Interestingly, neuronal presynaptic terminals become silent during excitotoxic insults, perhaps in a neuroprotective effort to avoid further damage, resulting in a period of abnormally reduced neuronal firing (Hogins et al., 2011). This progression from hypoactivity to hyperactivity has been shown in a controlled cortical impact model of TBI (Ping and Jin, 2016). Cortical deafferentation, another deprivation paradigm modeling TBI-induced epilepsy (Fröhlich et al., 2008), has been linked to homeostatic readjustment previously, with increased excitatory neuron intrinsic excitability and input resistance (Avramescu and Timofeev, 2008) as well as excitatory synapse scaling (González et al., 2015). Stimulation of cortex can provide relief from certain consequences of TBI, such as tinnitus, acquired epilepsy, and neuropathic pain (Chai et al., 2019). This suggests that the maladaptive compensatory plasticity observed *in vivo* and *in vitro* may have broader relevance to other pathological conditions giving rise to seizure disorders.

### Mechanisms of prolonged vs. short-term homeostatic compensation

Prior mechanistic studies of the homeostatic response to activity blockade have emphasized the importance of three coordinated mechanisms: a) a strengthening of excitatory synapses through postsynaptic upscaling, presynaptic enhanced release or both; b) a weakening of inhibitory synapses, also potentially occurring pre- and postsynaptically; and c) an increase in intrinsic excitability occurring either globally, or selectively involving the sensitivity, size and placement of the axon initial segment (Tien and Kerschensteiner, 2018; Turrigiano and Nelson, 2004; Wefelmeyer et al., 2016).

Surprisingly, we found limited support for the idea that persistent network hyperexcitability involved a lasting increase in the number or strength of excitatory synapses. Specifically, we found that excitatory synapse number decreased by 57% during the course of 10 days of deprivation followed by three days of recovery. A more intermediate value after 5 days of deprivation and three days of recovery was not significant. Prior studies of activity deprivation have attributed a similar progressive decrease in excitatory synapse density (Minerbi et al., 2009) to a failure to stabilize transient synapses (De Roo et al., 2008b, 2008a). Loss of excitatory synapses, while operating in the opposite direction expected to address an activity deficit, still could contribute to synchronicity if fewer synapses are unable to meet the demanding tuning requirements of a balanced circuit state (Atallah and Scanziani, 2009; Treviño, 2016).

There was no structural evidence of a lasting significant increase in synaptic size. A recent study in neocortical slice culture demonstrated that 5 days of deprivation without additional recovery period produced increases in synapse size evident both pre- and postsynaptically and apparent in both super resolution light microscopy and ultrastructurally (Wise, Escobedo-Lozoya et al. 2023). The difference between the present results and those obtained in the Wise study and in multiple other studies of deprivation-induced homeostatic plasticity (Barnes et al., 2017; Hobbiss et al., 2018; Mitra et al., 2011; Murthy et al., 2001; O’Brien et al., 1998; Turrigiano et al., 1998) suggests that this prominent feature of synaptic change may readjust rapidly as activity changes, and that deprivation-induced increases in synaptic size were likely reversed by elevated activity during the three day recovery period. Indeed, when TTX is applied to organotypic hippocampal cultures for a much shorter time (48 hours), synaptic scaling results in increased mEPSC amplitude acutely but presents a complete recovery when slices recover in media for one week (Koch et al., 2010). Because we did not obtain physiological measurements of spontaneous or evoked synaptic transmission, we cannot rule out the possibility of a lasting increase in the probability of presynaptic release. However, in all of the prior studies just mentioned, increased release was accompanied by increased presynaptic size, which was not seen here.

We furthermore did not observe evidence of a lasting decrease in the size of individual inhibitory synapses as reported previously in dissociated culture (Kilman et al., 2002; Swanwick et al., 2006). In fact, after ten days of activity blockade, followed by three days recovery, puncta of colocalized presynaptic VGAT showed a modest, but significant increase. Elevated extracellular potassium, which depolarizes cells and creates a hyperexcitable regime, is known to enlarge inhibitory synapses over 48 hours (Rannals & Kapur, 2011). During the overshoot period in our model, a similar hyperexcitability is induced. The increase we see in inhibitory presynapse size, therefore, could reflect an indirect transient change in response to the elevated activity of the recovery period and not a direct response to deprivation.

In contrast to the small increase in presynaptic inhibitory size, we also saw persistent and dramatic changes were evident in the number of presumed inhibitory synapses. Presumptive inhibitory synapses were reduced to half of their initial density by 5 days, and to a third of their initial density after 10 days of deprivation and in both cases these reductions were persistent since they were apparent after a three-day recovery period. Inhibitory synapse development has long been known to depend bidirectionally on circuit activity. Elevated activity increases the number of inhibitory synapses (Marty et al., 2000) as well as their size (Rannals and Kapur, 2011), and reduced activity decreases inhibitory synapse density (Chattopadhyaya et al., 2004) and/or size (Kilman et al., 2002). Epilepsy has also been linked to inhibitory synapse loss in humans (Alonso-Nanclares et al., 2011; Calcagnotto et al., 2005). Overall reduced inhibition likely contributed to the persistent epileptiform activity evident here after prolonged deprivation in slice culture.

Although both excitatory and inhibitory putative synapse density decreased, the loss was greater for inhibitory synapses (57% for excitatory synapses *vs*. 67% for inhibitory synapses). The balance between excitation and inhibition in neural circuits has been long understood as a functional requirement of healthy activity propagation (Chen et al., 2022; Sohal and Rubenstein, 2019; Zhou and Yu, 2018). This shift in propagation equilibrium is likely a contributor to the hypersynchronous circuit resulting from activity deprivation.

### Intrinsic excitability

In addition to a lasting reduction in inhibition, we also observed a lasting increase in intrinsic excitability. Layer 5 pyramidal cells exhibited a roughly twofold increase in input resistance and an elevated FI-slope after ten days of deprivation, and showed an intermediate, but nonsignificant level of firing after five days of TTX. Attempts to quantify the impact of deprivation prior to recovery were difficult because nearly all cells exhibited depolarization block (not shown), suggesting some degree of recovery of normal firing properties during the recovery period. Plasticity of intrinsic excitability in a homeostatic context was described alongside the early work on synaptic scaling (Desai et al., 1999), with excitability rising to produce greater firing rates following deprivation. Modeling studies have suggested that homeostatic intrinsic plasticity might contribute to seizures following brain injury and associated silencing (Houweling et al., 2005). Intrinsic excitability-based homeostatic plasticity is thought to be developmentally regulated (Wen and Turrigiano, 2021), which might explain in part how activity deprivation is so devastating during early postnatal development. In any case, although it is not surprising that neurons become more excitable after deprivation, we were surprised to find that these changes could persist after three days of recovery. These biophysical properties of the cell are very important to network function and the persistent shifts we observed in input resistance and firing output in pyramidal excitatory neurons push the cortical network towards persistent hyperexcitability.

## Conclusion

In conclusion, during protracted activity deprivation maladaptive compensatory changes in cortical circuits continue to accrue. Much as occurs with *in vivo* activity blockade, these changes are developmentally regulated and progressive and do not require Hebbian plasticity. Unlike *in vivo*, hyperactivity in cortical slice culture eventually recovers though this process takes many days. Mechanistically, we found that both inhibitory and excitatory synapses were lost, with a greater loss of inhibition, leading to an imbalanced circuit. In addition, changes in cellular excitability were surprisingly persistent and so likely also contribute to circuit hyperexcitability. Operationally, the slice culture preparation studied here may provide a useful testbed for future studies of pathophysiological mechanisms and potential treatments for seizure secondary to activity deprivation.

---

**Supplemental Figure 1:**
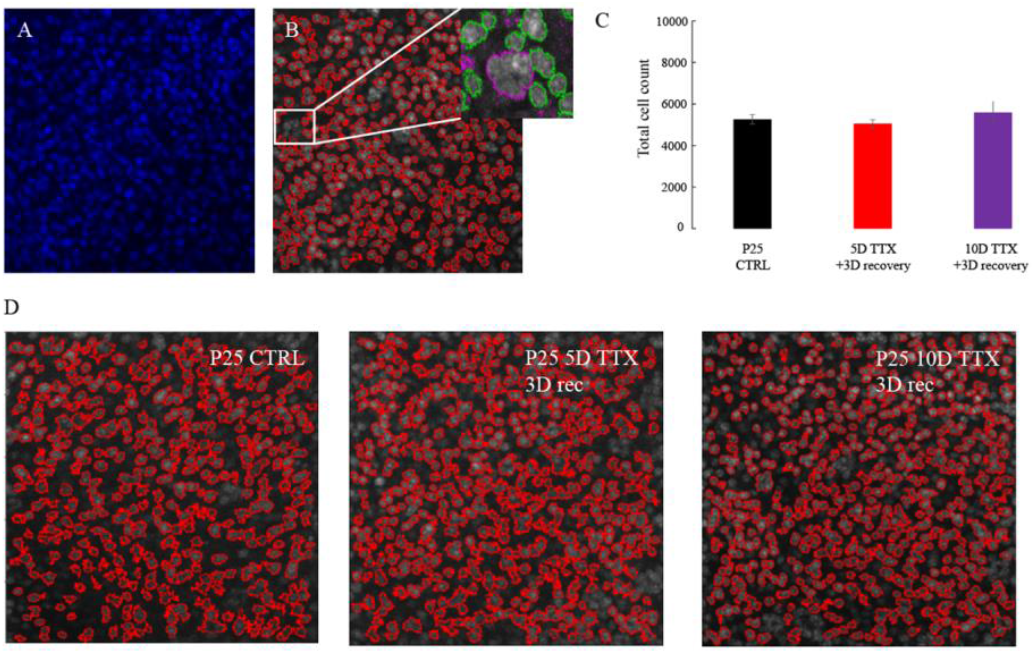
No significant cell death occurs with prolonged tetrodotoxin exposure. **A**. Example 10x image from Zeiss Airyscan of DAPI-stained nuclei (blue) in layer 5 of somatosensory cortex in 461 x 461 μm area. Full z extent was imaged for each stack, so the total volume differed in for each sample. **B**. CellProfiler calls (red outline) of good nuclei with inset showing examples of rejected blurred nuclei out of focus (purple outlines) and accepted crisp outlines of counted nuclei (green outlines). **C**. Bar graph of total nuclei count in our three conditions, from the same slices used in synaptic imaging analysis. **D**. Examples of accepted cell density (red outlines) from all three conditions.

